# Structural basis of pseudoGTPase-mediated protein-protein interactions

**DOI:** 10.1101/2024.10.30.620932

**Authors:** Bing Wang, Rui Yang, Chun Wan, Yuan Tian, Jingyi Wu, Sayantan Roy, Suzhao Li, Jingshi Shen, Qian Yin

## Abstract

GTPases regulate various cellular processes through conformational changes triggered by GTP or GDP binding. Recently, pseudoGTPases, the catalytically inactive counterparts of GTPases, have been identified across species from bacteria to human, although their functions and mechanisms remain unexplored. In this study, we demonstrate that the N-terminal region of the assembly chaperone AAGAB is a type i pseudoGTPase using biochemistry and X-ray crystallography. Furthermore, we discovered that the AAGAB pseudoGTPase domain (psGD) interacts with the σ subunits of AP1 and AP2 adaptor complexes, heterotetrameric complexes involved in clathrin-mediated membrane trafficking. AAGAB psGD engages the σ subunits via a unique interface distinct from the conventional GTPase interacting regions. Further biochemical and cell-based assays confirmed the crucial role of the newly identified interface in binding and membrane trafficking. Collectively, our results establish AAGAB pseudoGTPase domain as a critical protein-protein interaction module. These findings offer new insight into the structural basis and molecular mechanisms of pseudoGTPases.

**Highlights:**

- AAGAB N-terminal region is a pseudoGTPase
- AAGAB pseudoGTPase domain interacts with and stabilizes AP1 and AP2 σ subunits
- The crystal structure of AAGAB psGD:AP1σ3 is the first reported psGD complex structure, revealing a unique interface independent of guanine nucleotide regulation
- The psGD:AP1σ3 structure offers mechanistic insights into σ subunits stabilization and protection through adaptor complex assembly

## Introduction

GTPases are ubiquitous in all aspects of cellular processes, regulating cell signaling, transcription, translation, membrane trafficking, and cell motility. GTPases’ functions lie in their capability to adopt distinct conformations when bound to different nucleotides – “on” when bound to GTP and “off” when bound to GDP. The GTPase domain, or GTP-binding domain (GD), is usually composed of 160-180 residues and characterized by five signature GTPase motifs, G1-G5, that coordinate the guanine nucleoside, the phosphates, and the divalent ion essential for the GTP binding and hydrolysis cycle ^1–3^. The G2 and G3 motifs and surrounding regions are also known as the “switch I” and “switch II” regions, respectively. Switch I and II regions display substantial conformational changes based on bound nucleotides, enabling them to engage or disengage a variety of regulatory molecules or effector proteins to turn on or off cellular processes.

Recent studies have uncovered several noncanonical GTPases, from prokaryotes up to the human genome ^4^. They are similar to canonical GTPases in sequences and structures, yet they are incapable of hydrolyzing or binding GTP. Such noncanonical GTPases or domains are therefore named pseudoGTPases ^4^. A handful of pseudoGTPases are known to mediate kinetochore assembly, load cargo adaptor proteins to dynein heavy chain, or regulate cytoskeleton organization or other enzymatic activities ^4–8^. PseudoGTPases are widespread and implicated in diverse biological pathways, yet their functions and molecular mechanisms remain largely unknown.

In this work, we identified another pseudoGTPase domain (psGD) located at the N-terminus of AAGAB (alpha and gamma adaptin binding protein, also known as p34). AAGAB is the assembly chaperone for AP1 and AP2 adaptor complexes that are key to clathrin-mediated membrane trafficking ^9,10^. Assembly chaperones are proteins that interact with individual subunits and assembly intermediates during assembly of multi-subunit macromolecular complexes ^11^. While the roles of assembly chaperones are well recognized in the assembly of ribosomes, proteasomes, and nucleosomes, dedicated assembly chaperones have also been identified for SNARE complex assembly and disassembly ^12,13^. In AP1 and AP2 assembly, AAGAB forms ternary complexes with AP1γ and AP1σ subunits, or AP2α and AP2σ subunits, stabilizing the AP1γ:AP1σ and AP2α:APσ hemicomplexes ^9,10^. More recently, the chaperone function of AAGAB has been expanded to AP4 assembly ^14^. AAGAB-mediated assembly of the AP complex drives the biological process known as chaperone-assisted adaptor protein assembly (CAPA). Heterozygous mutations in the *AAGAB* gene cause the autosomal hereditary skin disease punctate palmoplantar kerotoderma type I (PPKP1, also known as Buschke-Fischer-Brauer) ^15,16^. The physiological importance of AAGAB is further highlighted by a recent article reporting a loss-of-function mutation in the *aagab* gene results in impaired swimming and early larval mortality in zebrafish ^17^.

AAGAB contains two conserved, structured regions: the N-terminal G domain, and the C-terminal domain ^18^. The C-terminal domain has been shown to mediate AAGAB self-dimerization and interaction with AP1γ and AP2α subunits ^18^. In comparison, the function of the highly conserved N-terminal G domain remains uncharacterized.

Here, we show that the N-terminal G domain (residues 2-177) is a pseudoGTPase domain (psGD). Through biochemistry and X-ray crystallography, we discovered that although the N-terminal G domain of AAGAB assumes a canonical GTPase fold, it lacks both GTPase activity and guanine nucleotide binding capacity, classifying it as a class i pseudoGTPase. Instead, AAGAB psGD interacts with and stabilizes the σ subunits of the AP1 and AP2 adaptor complexes. We then determined the crystal structure of the psGD in complex with AP1 σ subunit at the resolution of 1.7 Å. The high resolution psGD:AP1σ crystal structure reveals that AAGAB psGD grabs AP1σ subunit using two highly conserved loops in a pincer-like manner. The interface psGD utilizes to engage the σ subunit is distinct from the switch I and switch II regions widely used by small GTPases to engage GAPs, GEFs, or effector proteins. Further biochemical and cell-based assays confirmed the crucial role of the newly identified interface. Collectively, our results established the role of AAGAB psGD in chaperoning the assembly of AP complexes by interacting with, stabilizing, and protecting the σ subunits during AP assembly. Our results also revealed one of the functions of pseudoGTPase domains as a protein-protein interaction module. Thirdly, the distinct interface used by AAGAB psGD points to an interface independent of guanine nucleotide binding, hinting at an independently evolved path and may assist future protein engineering.

## Results

### AAGAB contains a pseudoGTPase domain (psGD) at its N-terminus

The N-terminal conserved region of human AAGAB is annotated as a G domain (GD). As we scrutinize its sequence, however, we notice that several key G motif residues are missing or profoundly altered in AAGAB GD, particularly in G1 and G3 (Figure 1a and Suppl. Figure 1a). As a first step to determine its function, we overexpressed and purified the AAGAB N-terminal region (residues 2-177) from *E. coli*. AAGAB N-terminal region existed as a monomer in solution, as evaluated by size-exclusion chromatography (SEC) elution position and confirmed by SEC-coupled multi-angle light scattering (SEC-MALS) (Suppl. Figure 1b) ^18^. Confirming our doubt, AAGAB GD displayed no GTPase activity when the positive control guanylate-binding proteins 2 (GBP2) steadily hydrolyzed GTP (Figure 1b). Furthermore, AAGAB GD did not bind to fluorescently labeled nonhydrolyzable GTP mimic or GDP even at the concentration of 40 μM (Figure 1c, Suppl. Figure 1c). These data suggest that AAGAB does not bind or hydrolyze GTP, hinting at the classification as a pseudoGTPase. To further test this possibility, we resorted to crystallography to determine whether the putative G domain truly adopts a GTPase fold. Despite extensive effort, the wild type G domain did not produce any crystals. However, GD protein bearing a disease-related mutation, E144K ^19^, successfully crystallized and diffracted to high resolution. The GD^E144K^ mutant behaves almost identical to the wild type protein in all assays tested (Suppl. Figure 1d and see below). We believe it represents the characteristics of the wild type protein. We determined the crystal structure of AAGAB GD^E144K^ using molecular replacement and refined the structure to 1.78 Å (Figure 1d, Suppl. Figure 1e, Table S1). The two GD molecules in each asymmetric unit superpose well with each other, with a root mean square deviation (RMSD) of 0.341 Å across 142 α carbon atoms (Cαs), the only major difference coming from the loop connecting α1 to β2 (Suppl. Figure 1e). Therefore, we will use molecule A for all structural descriptions below. The structure of AAGAB N-terminal G domain indeed reveals a small GTPase fold, with the topology of one central six-stranded β sheet sandwiched by five α helices, three of which (α2/α2’, α3, and α4) on one side and the other two (α1, α5) on the opposite side (Figure 1d-e, Suppl. Figure 1e-f). The central β sheet is in a mixed arrangement where five parallel strands (β1, β3 - β6) are flanked at one side by the antiparallel peripheral β2. The topology of AAGAB G domain is the same as well-characterized small GTPases such as HRas ^20^ (Figure 1e, Suppl. Figure 1f). Although the overall framework of AAGAB GD closely resembles that of HRas, the loops connecting the secondary structures adopt vastly different conformations. In particular, the loop connecting β1 and α1, equivalent to the G1 motif in small GTPases, clashes with guanine nucleotide binding. The loop connecting α1 to β2, equivalent to the G2 motif/ “switch I” region in small GTPases, moves far away from the nucleotide binding site. At the other end of the nucleotide binding pocket, the loops connecting β5 to α4 and β6 to α5, counterparts of G4 and G5 motifs in small GTPases, also display large-scale movements to the point that they no longer support guanine binding ^20,21^ (Figure 1e, Suppl. Figure 1f). Consistently, we did not observe any GTP or GDP electron density in the crystal structure of AAGAB psGD. Collectively, our biochemical and structural results classify AAGAB N-terminal region as a class i pseudoGTPase: incompetent of GTP binding and hydrolysis ^4^. Hereafter, we will use pseudoGTPase domain (psGD) to describe the N-terminal region of AAGAB.

**Figure 1.**
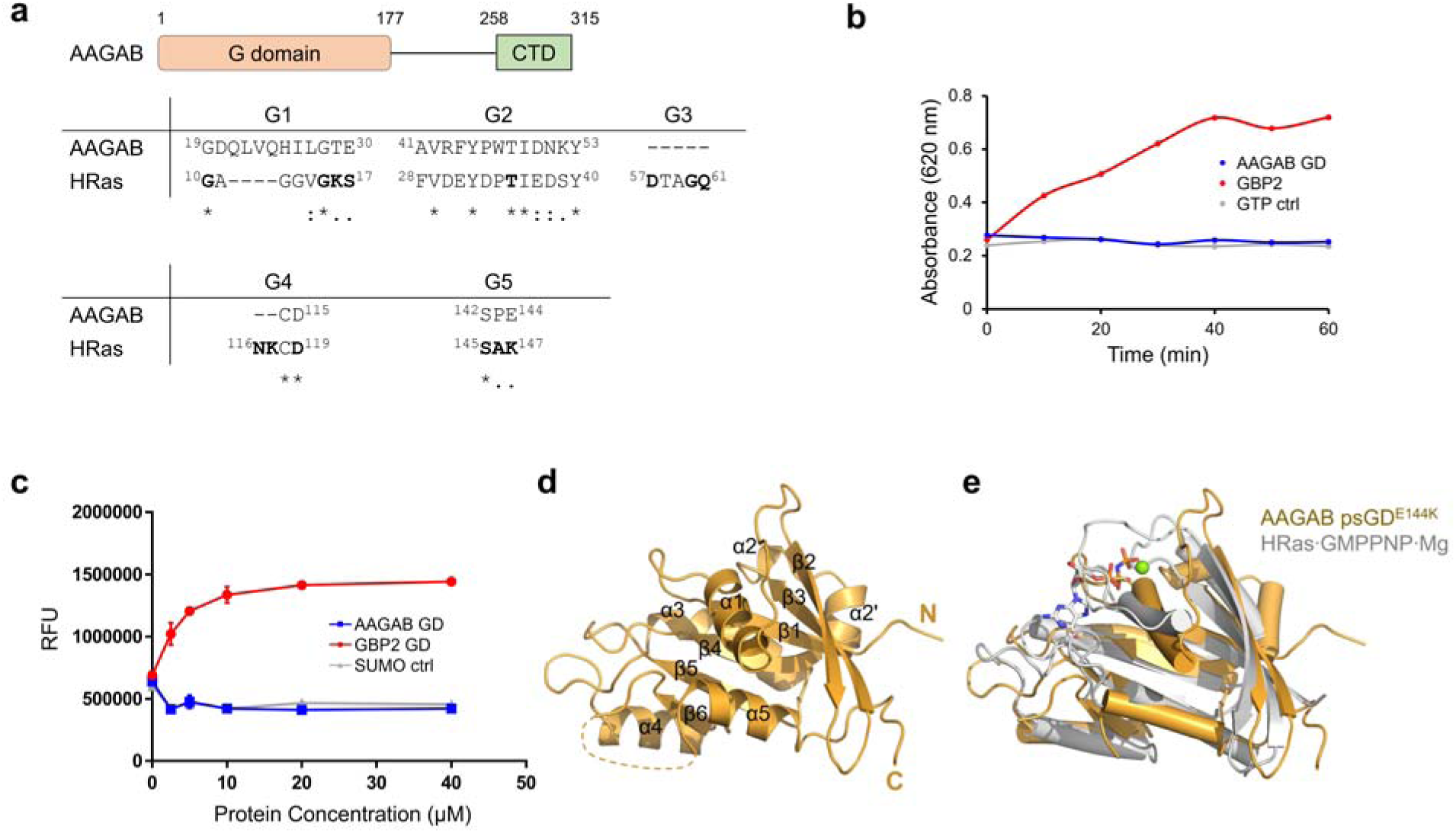
AAGAB N-terminal region is a pseudoGTPase domain. (a) Schematic diagram of AAGAB (upper) and sequence alignment of G motif residues between AAGAB N-terminal region and HRas (lower). Residue numbers are labeled as superscripts. Consensus G motif residues in HRas are shown in bold. (b) GTPase activity of AAGAB GD as measured by malachite green assay. Full-length GBP2 and GTP are included as positive and negative controls. (c) AAGAB GD, GBP2 GD, and SUMO are mixed with mant-GTPγS and changes in mant-GTPγS fluorescence are monitored. (d) Crystal structure of AAGAB GD^E144K^. The N- and C-termini and secondary structure elements are labeled. (e) Structure superposition of AAGAB psGD (gold) with HRas·GMPPNP·Mg (gray, PDB ID 5P21).

AAGAB is evolutionarily conserved in eukaryotic organisms. A yeast protein Irc6p has been proposed as AAGAB homolog based on the experiment that human AAGAB can rescue growth defect in yeast irc6Δ cells ^22,23^. To explore the molecular basis for the conserved and cross-species function, we compared the structures of the pseudoGTPase domains of AAGAB and Irc6p (Suppl. Figure 1g and 1h) ^23^. The RMSD is 2.143 Å over 120 Cα atoms. The central β sheets superpose well with each other, as well as the α2, α2’, α3, and α4 helices on one side of the β sheet. α1 and α5 on the other side, however, display substantial movements (Suppl. Figure. 1h). α1 in AAGAB swivels ∼ 28° compared to α1 in Irc6p. The loops connecting α1 and β2, β5 and α4, and β6 and α5, and the β3-α2’ junction also display large-scale movements. To summarize, AAGAB psGD and Irc6p GD share highly similar framework but very dynamic loops.

### AAGAB pseudoGTPase domain is sufficient and necessary to interact with and stabilize AP1 and AP2 σ subunits

Small GTPases are widely used in cell signaling pathways as “switches”. Depending on their guanine nucleotide binding states, small GTPases can associate or dissociate from a wide range of effector molecules, thus turning “on” or “off” signaling. AAGAB is reported to bind to AP2α and AP2σ subunits and form a stable AAGAB:AP2α:AP2σ ternary complex ^9^. As the AAGAB C-terminal region is responsible for AP2α (or AP1γ subunit) interaction using its C-terminal helical domain ^18^, we are interested in exploring whether the N-terminal psGD can bind the σ subunits.

We coexpressed N-terminally GST-tagged AAGAB psGD (GST-psGD) and C-terminally His_6_-tagged AP1σ3 (AP1σ3-His_6_) in *E. coli* and purified the cell lysates with tandem Ni and GST affinity chromatography. In the Ni-NTA elution fraction we clearly identified two major bands corresponding to GST-psGD and AP1σ3-His_6_ (Figure 2a, lane 2), suggesting direct interaction between the two proteins. The Ni-NTA elute fraction was further purified by a second round GST affinity chromatography. Both GST- psGD and AP1σ3-His_6_ bound to glutathione beads, confirming the direct interaction between psGD and AP1σ3 (Figure 2a, lane 4).

**Figure 2.**
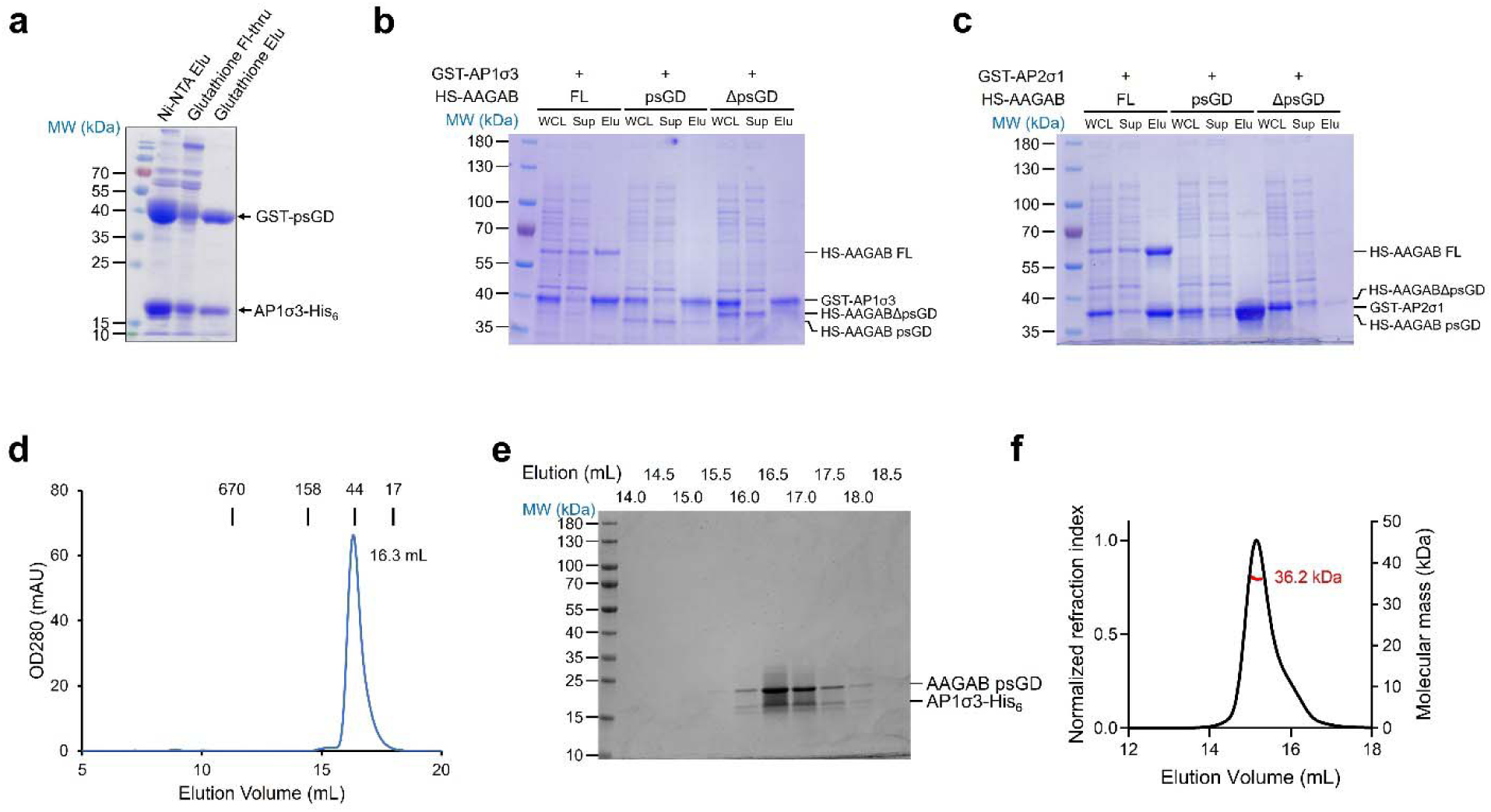
AAGAB psGD interacts and stabilizes AP1 and AP2 sigma subunits. (a) GST-psGD and AP1σ3-His_6_ are co-expressed in *E. coli*. The cell lysate is subjected to Ni-NTA pull down followed by glutathione pull down. AP1σ3-His_6_ pulls down GST- psGD and vice versa. Elu: elution; Fl-thru: flow-through. (b) GST pull-down SDS-PAGE gel of GST-AP1σ3 co-expressed with His_6_-SUMO (HS) tagged full-length (FL) AAGAB, AAGAB psGD (2-177), or AAGABΔpsGD (178-315). WCL: whole cell lysate; Sup: supernatant; Elu: glutathione elution. (c) GST pull-down SDS-PAGE gel of GST-AP2σ1 co-expressed with His_6_-SUMO (HS) tagged full-length (FL) AAGAB, AAGAB psGD (2- 177), or AAGABΔpsGD (178-315). WCL: whole cell lysate; Sup: supernatant; Elu: glutathione elution. (d) SEC profile of AAGAB psGD:AP1σ3-His_6_ binary complex. Protein standards with known molecular weights are marked at the top. (e) SDS-PAGE of the SEC fractions in (d). (f) SEC coupled multi-angle light scattering (SEC-MALS) profile of AAGAB psGD:AP1σ3-His_6_ binary complex.

We further validated the interactions by coexpressing GST-tagged AP1σ3 (GST- AP1σ3) with N-terminal His_6_-SUMO (HS)-tagged full-length (FL) AAGAB (residues 2- 315), AAGAB psGD (residues 2-177), and AAGABΔpsGD (residues 178-315) in *E. coli*. GST-AP1σ3 was able to pull down FL AAGAB and psGD, but not AAGABΔpsGD (Figure 2b), corroborating the conclusion that AAGAB psGD is sufficient and necessary to interact with AP1σ3. As AAGAB has been shown to chaperone the assembly of both AP1 and AP2 complexes ^9,10,18^, we examined whether psGD interacts with AP2σ subunit. Indeed, GST-AP2σ1 was able to pull down HS-tagged FL AAGAB and psGD, but not AAGABΔpsGD in the bacterial coexpression system, mirroring the results of AP1σ3 (Figure 2c).

When overexpressed in *E. coli* by itself, a small amount of the AP1 or AP2 σ subunit was soluble. Yet this small amount of σ subunit was mostly misfolded, as demonstrated by the elution position of GST-AP1σ3 by size-exclusion chromatography (SEC, Suppl. Figure 2a and 2b). To confirm that AAGAB aids the solubility and folding of σ subunits, we characterized the coexpressed and co-purified AAGAB psGD:AP1σ3- His_6_ complex by SEC (Figure 2d and 2e). The binary complex, in contrast, eluted as a well folded, soluble entity at ∼ 16.3 mL, corresponding to a molecular weight of ∼ 45.9 kDa. Both the molecular weight and the SDS-PAGE of SEC peak fractions suggest a 1:1 stoichiometry between AAGAB psGD and AP1σ3 (Figure 2d and 2e). Furthermore, size-exclusion chromatography coupled multi-angle light scattering (SEC-MALS) revealed a molecular mass of 36.2 kDa for the binary complex (Figure 2f), consistent with calculated molecular mass of 38.8 kDa for the 1:1 AAGAB psGD:AP1σ3 complex.

### psGD:AP1σ3 crystal structure reveals a unique interface

To further understand the molecular basis of the interactions between AAGAB psGD and the σ subunits, we determined the crystal structure of wild type AAGAB psGD in complex with AP1σ3 at the resolution of 1.68 Å (Figure 3a, Suppl. Figure 3a). The complex structure was determined by molecular replacement using AP1σ1 structure as a searching model (PDB ID: 1W63, chain U) ^24^.

**Figure 3.**
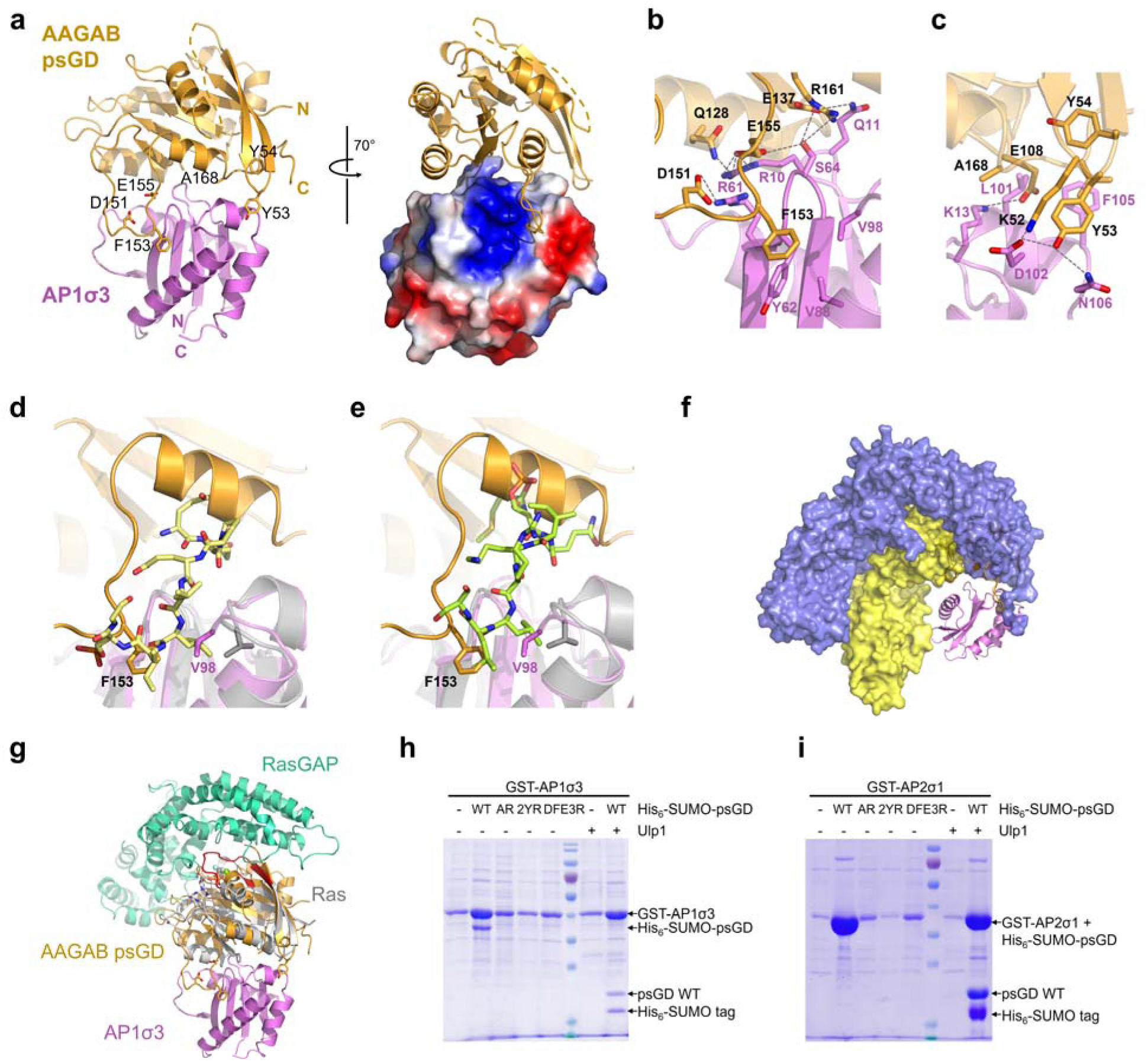
AAGAB psGD utilizes a unique interface to interact with sigma subunits. (a) Cartoon and surface representation of AAGAB psGD:AP1σ3 complex structure with psGD in gold and AP1σ3 in violet. N- and C- termini of both proteins are labeled. Key interfacial residues on psGD are shown as sticks and labeled. In the right panel, the complex is rotated by 70 degrees and AP1σ3 is shown as electrostatic surface. Residues missing electron densities are represented by the dashed line. (b) Zoomed-in view of the psGD D151/F153/E155 interaction with AP1σ3. (c) Zoomed-in view of psGD A168 and K52/Y53 interactions with AP1σ3. (d) Superposition of the AAGAB psGD:AP1σ3 with the AP1σ:pSTING complex (PDB ID 7R4H). The AP1σ is colored gray and the pSTING peptide is shown as stick in pale yellow. F153 in AAGAB psGD and V98 in AP1σ3 are shown as sticks and colored orange and violet, respectively. (e) Superposition of the AAGAB psGD:AP1σ3 with the AP2core in complex with two cargo peptides (PDB ID 6QH6). The AP2σ is colored gray and the peptide is shown as stick in pale green. F153 in AAGAB psGD and V98 in AP1σ3 are shown as sticks and colored orange and violet, respectively. (f) Superposition of the AAGAB psGD:AP1σ3 with AP1 core in closed form (PDB ID 1W63). The AP1β and AP1μ subunits are shown as surface and colored in blue and yellow. (g) Superposition of psGD: AP1σ3 with HRas: RasGAP structure (PDB ID 1WQ1). AAGAB psGD and AP1σ3 are colored gold and violet as in (a). HRas and RasGAP are colored in gray and greencyan, respectively. The switch I and II regions of HRas are highlighted in red. (h) GST pull-down gel of GST-AP1σ3 co- expressed with His_6_-SUMO tagged WT or mutant AAGAB psGD. AR: A168R; 2YR: Y53R/Y54R; DEF3R: D151R/F153R/E155R. Ulp1 protease treatment is used to confirm the identity of His_6_-SUMO-AAGAB psGD^WT^. (i) GST pull-down gel of GST-AP2σ1 co- expressed with His_6_-SUMO tagged WT or mutant AAGAB psGD. AR: A168R; 2YR: Y53R/Y54R; DEF3R: D151R/F153R/E155R. Ulp1 protease treatment is used to confirm the identity of His_6_-SUMO-AAGAB psGD^WT^. Note that the molecular weights of GST- AP2σ1 and His_6_-SUMO-AAGAB psGD are too close for separation on SDS-PAGE.

In the crystal structure AAGAB psGD forms a 1:1 complex with AP1σ3. The overall structure of psGD in the complex is similar to the free psGD^E144K^ structure with an RMSD of 0.577 Å for 149 aligned Cα atoms. The secondary structures of the psGD domains are almost identical. The differences lie in the flexible S1 loop, the loop connecting α1 to β2, and the loop connecting β6 and α5 (Suppl. Figure 3b). The S1 loop is missing in the psGD:AP1σ3 structure, consistent with its flexible nature even in isolated psGD structures (Suppl. Figure 3b). On the other hand, the β6-α5 loop, missing in the isolated psGD structures, is well resolved and makes intimate and extensive interactions with AP1σ3 (Figure 3a and 3b). In the complex, AP1σ3 folds with a central five-stranded antiparallel β sheet sandwiched by two layers of helices (Figure 3a), a typical fold of longin and roadblock domains (LD/RDs) which is an ubiquitous and conserved scaffold for small GTPases in various biological processes across species ^25–29^. The overall structure of AP1σ3 in our complex is also similar to the AP1σ1 structure in the AP1 core structure ^24^. However, the electron density for the last alpha helix (residues Q124-R154) in AP1σ3 is missing. The equivalent alpha helix in the AP1 core structure has minimum interaction with other subunits, suggesting a certain degree of structure flexibility.

The psGD:AP1σ3 interface buries an area of 994 Å^2^ as calculated by the PDBePISA server ^30^. In addition, the complex formation significance score is 1.000, implying the interface plays an essential role in complex formation. AAGAB psGD envelopes one end of the β sheet in AP1σ3, tightly wrapping around the β1-β2 and β4- β5 loops and making additional contacts with the surrounding alpha helices (Figure 3a, Suppl. Figure 3a). The intermolecular interaction network of psGD:AP1σ3 is mediated mostly by charged or polar residues with moderate contributions from hydrophobic residues (Figure 3a-c). At one side of the AP1σ3 β sheet, two loops on AAGAB psGD protrude prominently and grab AP1σ3 like a pincer (Figure 3a, Suppl. Figure 3a). The longer “jaw” of the pincer is composed of the β6-α5 “substrate/σ subunit binding loop” (SBL) that is unique in AAGAB psGD (Figure 3a-b, Suppl. Figure 3a). Notably, this loop region is flexible in isolated AAGAB psGD structure and only becomes ordered when binding to AP1σ3. D151, F153, and E155 of AAGAB form the nucleus of this interface. In the center, the aromatic side chain of F153 fits into a hydrophobic pocket lined by Y62, V88, and V98 in AP1σ3, and the mainchain carbonyl oxygen of F153 forms hydrogen bonds with both the sidechain of R61 and mainchain nitrogen of A63 in AP1σ3 (Figure 3b). D151 and E155 form salt bridges with R61 and R10 in AP1σ3, respectively (Figure 3b). This interface is further bolstered by additional interactions including hydrogen bonding between Q128 and AP1σ3 R10, E137 and AP1σ3 Q11, E155 and AP1σ3 S64, and R161 and AP1σ3 S64 (Figure 3b). The shorter “jaw” of the pincer consists of the short β2-β3 loop that is highly conserved in AAGAB proteins across species ^18^. K52 and Y53 form the core of this interface. K52 interacts with AP1σ3 D102 via a salt bridge, while Y53 simultaneously hydrogen bonds with both D102 and N106 in AP1σ3 (Figure 3c). In addition, the aromatic moiety of AAGAB Y53 fits snugly in a pocket lined by AP1σ3 F105 and L101 (Figure 3c). Both hydrophilic and hydrophobic interactions connect the two jaws, essentially forming a continuous interface between AAGAB psGD and AP1σ3. For example, AAGAB E108 forms a salt bridge with K13 in AP1σ3 β1-β2 loop, and AAGAB A168 forms hydrophobic interactions with AP1σ3 L101 and L104 (Figure 3c). Additional interactions can be found at the other side of the AP1σ3 β sheet. AAGAB psGD α4 helix runs antiparallelly to the C-terminal end of AP1σ3 H1 helix, with AAGAB I132 contributing hydrophobic interactions with AP1σ3 L40 and S41 (Suppl. Figure 3c). The interfacial residues in AP1σ3 are highly conserved in AP1, AP2, and AP4 σ subunits (Suppl. Figure 3d), all of which depend on AAGAB for assembly ^9,10,14^.

One major sorting motif that links cargo proteins to AP1 or AP2 complexes is the dileucine-based motif [D/E]XXXL[L/I/M], where X is any amino acid ^31^. The dileucine motif binds to the σ subunit in an extended conformation, with the two leucines fitting in two adjacent hydrophobic pockets ^32–34^. We noticed that two of the hydrophobic residues lining the cargo peptide binding pockets, V88 and V98, participate in AAGAB psGD interaction as well (Figure 3b). Superposing our AAGAB psGD:AP1σ3 structure with AP1 or AP2 σ subunit loaded with cargo peptides immediately reveals steric clashes between the AAGAB psGD and the cargo peptides ^33,34^ (Figure 3d-e). α5 in psGD directly caps one end of the cargo peptide-binding grove, blocking the entry of the cargo peptide. More interestingly, at the dileucine binding site, V98 in AP1σ3 shifts ∼ 2.6 Å toward AAGAB psGD, occluding the L0 binding site. AAGAB F153 points toward AP1σ3, occupying exactly the same position as the L(+1) residue in the dileucine motif (Figure 3d). Together, AAGAB F153 and AP1σ3 V98 preclude cargo peptide binding at its C-terminus. Superposition with AP2σ reveals almost identical steric clashes (Figure 3e), suggesting AAGAB psGD blocks dileucine-based motif binding to AP2 as well.

Comparing psGD:AP1σ3 structure with fully assembled AP1 and AP2 core structures also reveals that psGD is incompatible with the β and μ subunits in the assembled core (Figure 3f), implying β and/or μ subunit can physically displace AAGAB to complete AP1 and AP2 assembly.

Although several pseudoGTPase structures in different biological processes have been reported ^5–8^, to our knowledge, there are no complex structures reported in relation to the class i pseudoGTPase lacking both nucleotide-binding and catalytic activity. To unravel its structural mechanisms, we compared the complex interface with representative small GTPase complex structures. Intriguingly, we found that AAGAB psGD uses a distinct interface for binding its partner. Typical small GTPases use the nucleotide binding pocket, the switch I and II regions, and surrounding regions to bind GTPase activating proteins (GAPs), guanine nucleotide exchange factors (GEFs), and effectors ^35–41^ (Figure 3g, Suppl. Figure 3e). Although the interfaces show slight differences in combination of structural elements, the location of the interface is very invariable ^35,36,41^ (Suppl. Figure 3e). Strikingly, AAGAB psGD uses a completely different surface, the surface on the opposite side of the nucleotide binding pocket, for engaging the AP1σ3 subunit (Figure 3a and 3g, Suppl. Figure 3e).

The AAGAB residues responsible for AP1σ3 interaction are highly conserved all the way down to yeast (Suppl. Figure 1g) ^18^, suggesting they are crucial in AAGAB function. We experimentally validate the interfacial residues by pull-down assay and mutagenesis. When GST-AP1σ3 was coexpressed with wild type (WT) His_6_-SUMO- psGD or mutants in *E. coli*, glutathione affinity purification was able to pull down both GST-AP1σ3 and WT His_6_-SUMO-psGD (Figure 3h, lane 2). The identity of the His_6_- SUMO-psGD band was further confirmed by Ulp1 treatment, which separates His_6_- SUMO from the psGD. In contrast, the AAGAB psGD A168R (AR), Y53R/Y54R (2YR), and D151R/F153R/E155R (DFE3R) mutants cannot be pulled down by GST-AP1σ3 (Figure 3h, lanes 3-5), confirming the crucial role of these residues in mediating AAGAB psGD:AP1σ3 interaction. Reciprocal pull down using Ni-affinity purification confirmed the interaction between AP1σ3 and WT psGD but not the mutants (Suppl. Figure 3f).

We noticed that when coexpressed with WT AAGAB psGD, the expression level of AP1σ3 increased substantially (Figure 3h), suggesting a role of AAGAB psGD in stabilizing AP1σ3. Similarly, we observed that GST-AP2σ1 can pull down coexpressed WT His_6_-SUMO-psGD, but not the AR, 2YR, or DFE3R mutants (Figure 3i and Suppl. Figure 3g), indicating AAGAB psGD adopts the same surface to interact with and stabilize AP2σ1.

### Disease-related mutations in psGD modestly reduces psGD stability

About 30 *AAGAB* mutations have been reported in patients with type I punctate palmoplantar keratoderma (PPKP1), a skin disease with the manifestation of skin thickening and pain on hands and feet ^15,16,18,19,42,43^. Most mutations are frameshift and nonsense mutations that lead to the loss of the C-terminal region of AAGAB protein.

However, two missense mutations, E144K and V139I, are located in the psGD region ^19,43^, prompting us to investigate their deficiencies. We generated E144K and V139I in the psGD protein. Both mutants expressed well and eluted at the same position as the wild type psGD protein. Coexpression of GST-AP1σ3 with N-terminally SUMO-tagged psGD WT, E144K, or V139I and subsequent glutathione affinity pull down experiment showed both E144K and V139I mutants interact with AP1σ3 (Suppl. Figure 4a).

Coexpression with GST-AP2σ1 yielded the same pull down results (Suppl. Figure 4b). We concluded that the E144K and V139I mutations do not disrupt the interaction between the psGD domain and the σ subunits. While E144 is on the psGD surface, V139 is partially buried. We then examined whether the mutations affect psGD stability. Using the Thermal Shift Dye as a reporter, we monitored the thermal stability for WT psGD, E144K, and V139I mutants (Suppl. Figure 4c). Compared to WT psGD that has a melting temperature (*T*m) of ∼ 61.1°C, the E144K mutant displayed a slight lower *T*m of ∼ 60.4°C, while V139I is modestly lower, with a *T*m of 59.4°C, 1.7°C lower than that of the WT (Suppl. Figure 4d). The results are consistent with the psGD structure in which the E144 residue is exposed and V139 is partially buried. Perturbation of the hydrophobic core of psGD by substituting a larger leucine sidechain for the small valine likely destabilizes psGD, leading to a reduced *T*m.

### psGD:σ interaction is crucial for σ subunit stability and membrane trafficking in the cell

Previously we have reported that AAGAB C-terminal domain is crucial for the stability of AP1γ or AP2α subunit and consequently AP1- and AP2-mediated clathrin-dependent membrane trafficking ^18^. To examine the physiological relevance of our structural and biochemical findings on AAGAB psGD, we assessed the effects of psGD mutations in cells. For cellular studies, we constructed C-terminally HA-tagged AP1σ3 and N-terminally 3xFlag-tagged AAGAB wild type (WT), Y53R/Y54R double mutant (2YR), and D151R/F153R/E155R/A168R/V171R quintuple mutant (DFEAV5R) (Figure 4a). 3xFlag- AAGAB proteins were transiently expressed in *AAGAB* KO HeLa cells with an empty vector or the plasmid encoding HA-tagged AP1σ3. Co-immunoprecipitation with anti-Flag antibody followed by anti-HA antibody clearly showed WT AAGAB, but not the psGD mutants, was able to pull down AP1σ3 subunit (Figure 4b, Suppl. Figure 5), consistent with in vitro biochemical observations (Figure 3h-i). The expression level of AP1σ3 was also lower when coexpressed with the psGD mutants (Figure 4b). Both AP1- and AP2-mediated membrane trafficking events are disrupted in *AAGAB* KO HeLa cells, leading to accumulation of transferrin receptor on the plasma membrane ^9,10,18^ (Figure 4c-d). Expression of WT AAGAB in *AAGAB* KO HeLa cells was able to restore surface TfR level. In contrast, the expression of psGD mutant 2YR and DFEAV5R did not restore the surface levels of TfR (Figure 4c-d), suggesting that disruption of psGD:σ interaction impairs membrane trafficking. We further examined the effect of psGD mutation on the stability of endogenous AP1 and AP2 subunits. Consistent with previous reports, expression of WT AAGAB protein in *AAGAB* KO HeLa cells greatly elevated the proteins levels of endogenous AP1γ, AP1σ, AP2α, and AP2σ subunits (Figure 4e-f).

**Figure 4.**
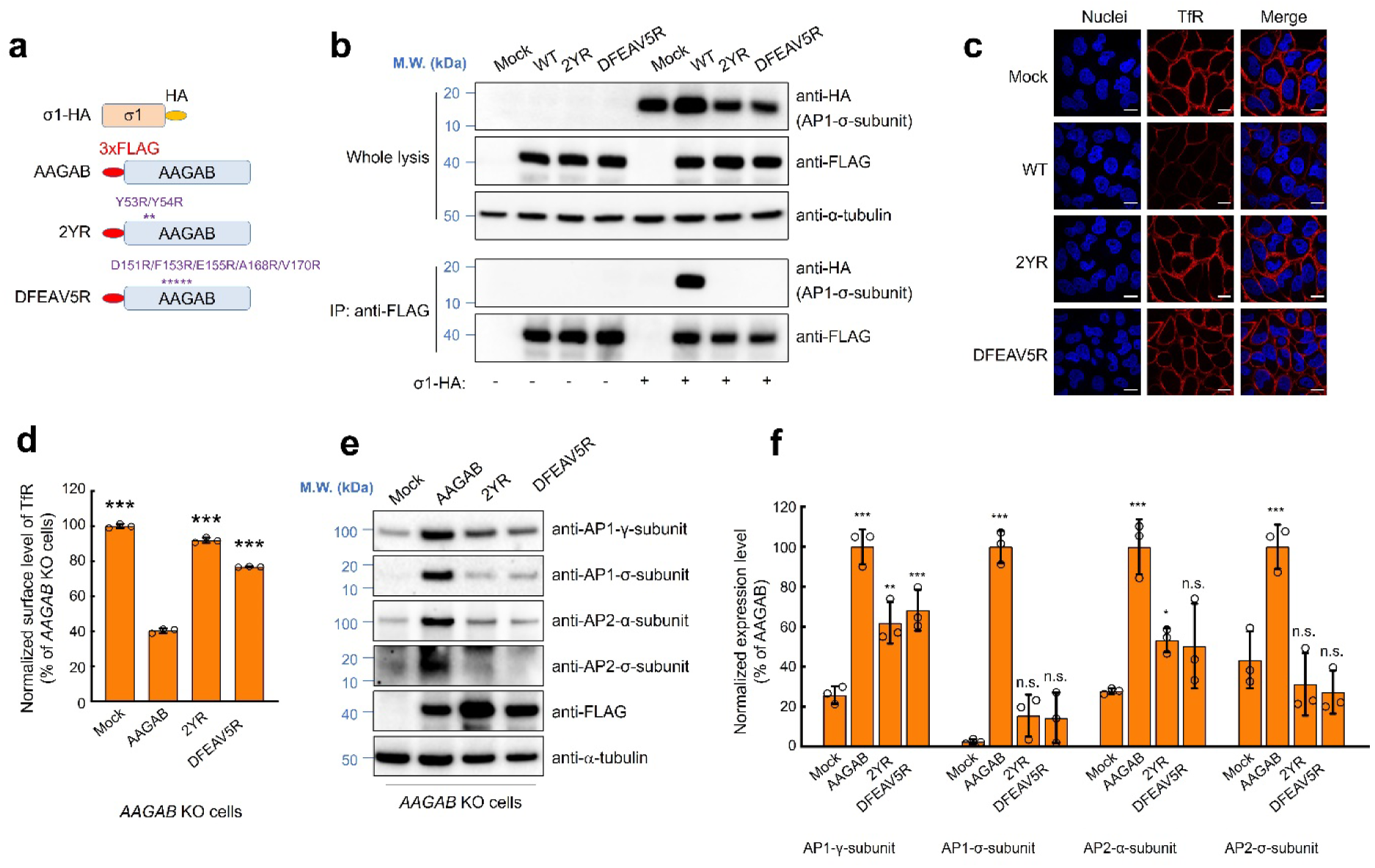
Disrupting psGD:σ interaction reduces σ subunit level in cells and compromises membrane trafficking. (a) Diagrams of HA-tagged full-length AP1 σ subunit and 3xFLAG-tagged WT or mutant AAGAB used in immunoprecipitation (IP) experiments. (b) Representative immunoblots showing the interaction of 3xFLAG-tagged WT or mutant AAGAB with HA-tagged AP1 σ subunit. The 3xFLAG-AAGAB proteins were transiently expressed in *AAGAB* KO HeLa cells with an empty vector (control) or plasmids encoding the HA-tagged AP1 σ subunit. The 3xFLAG-AAGAB proteins were immunoprecipitated from cell lysates using anti-FLAG antibodies, and the presence of 3xFLAG-AAGAB and HA-tagged AP1 σ subunit in the immunoprecipitants was detected using anti-FLAG and anti-HA antibodies, respectively. (c) Representative confocal microscopy images from three experiments showing surface levels of TfR in *AAGAB* KO HeLa cells expressing WT or mutant AAGAB. Surface TfR levels of non-permeabilized cells were labeled using anti-TfR antibodies and Alexa Fluor 568-conjugated secondary antibodies (red). Nuclei were stained with Hoechst 33342 (blue). Images were captured using a 100× oil immersion objective on a Nikon A1 Laser Scanning confocal microscope. Scale bars: 20 μm. (d) Flow cytometry measurements of the surface levels of TfR in *AAGAB* KO HeLa cells expressing the indicated proteins. Cells were disassociated by Accutase and stained with monoclonal anti-TfR antibodies and APC-conjugated secondary antibodies. APC fluorescence measurements of ∼5000 cells were collected on a CyAn ADP analyzer. Mean APC fluorescence was normalized to the control sample in which an empty vector was transfected into *AAGAB* KO cells. Data are presented as mean±s.d., n=3. ***P<0.001. P values were calculated using one-way ANOVA and Dunnett’s multiple comparison test. (e) Representative immunoblots showing the expression of the indicated proteins in *AAGAB* KO HeLa cell expressing WT or mutant AAGAB. (f) The relative expression levels of indicated proteins were normalized to WT AAGAB samples. Data are presented as mean±s.d., n=3. ***P<0.001, **P<0.01, *P<0.05, n.s., P>0.05. P values were calculated using one-way ANOVA and Dunnett’s multiple comparison test. The representative immunoblots are shown in (e). Mock: cells transfected with an empty vector.

Expression of psGD mutants 2YR and DFEAV5R, however, had minimum effect in elevating AP1σ or AP2σ protein expression in *AAGAB* KO cells (Figure 4e-f), underlining the critical role of psGD in stabilizing both σ subunits. In contrast, expression of AAGAB 2YR and DFEAV5R mutants moderately restored AP1γ and AP2α expression (Figure 4e-f), implying psGD is not involved in maintenance of AP1γ and AP2α stability. Together, these results are consistent with our in vitro data and demonstrate that psGD mediates the binding of AAGAB to AP1σ and AP2σ subunits in the cell.

## Discussion

We determined the crystal structure of AAGAB pseudoGTPase domain (psGD). Compared with canonical small GTPases, two features stand out in AAGAB psGD. The first is the α1-β2 “S1” loop that is equivalent to the G2 motif in canonical small GTPases. The G2 motif in small GTPases contacts with γ-phosphate and the Mg^2+^ ion, therefore stabilizing guanine nucleotide binding ^1^. The G2 motif is also known as the switch I region due to its highly dynamic nature in GTP- or GDP- binding states. In AAGAB psGD crystal structures, the equivalent S1 loop is also highly dynamic but often stays away from the canonical guanine nucleotide binding pocket (Figure 1d-e). The behavior of the S1 loop is in accordance with the lack of guanine nucleotide binding of AAGAB psGD.

On comparing the structures of psGD of AAGAB and Irc6p with canonical small GTPases, we find a unique feature in psGD, an extended loop connecting β6 and α5 (Figure 3a, 3b, Suppl. Figure 1a, 1f-1h). This loop is equivalent to the loop that houses the G5 motif ([C/S]A[K/L/T]) in canonical small GTPases ^1^ (Figure 1a, Suppl. Figure 1a). The consensus G5 motif is used by GTPases to specifically coordinate the guanine base not other nucleotides. The typical short G5 loop, together with the G4 motif, creates a closed pocket to wrap around the guanine base. In contrast, the loop in AAGAB is much longer than a typical G5 loop and sways away from the guanine base binding pocket (Suppl. Figure 1f and 1h). Both the lack of G5 consensus residues and the orientation of the β6-α5 loop are consistent with the absence of nucleotide binding capacity in AAGAB psGD. Instead, the β6-α5 loop acquires a new function in interaction with the AP1 and AP2 σ subunits, so we named it “substrate/σ subunit binding loop” (SBL) (Figure 3a-b, 3h-i). The length and amino acid composition of the SBL loop is conserved from yeast to human, hinting at an ancient functional role.

Fully assembled AP1 and AP2 core complexes adopt a “closed” conformation, in which the cargo peptide binding sites are protected. Membrane binding induces an “open” conformation to expose the cargo binding sites ^31^. However, it is unclear how the cargo binding site on σ subunits is shielded from untimely cargo binding during protein synthesis and complex assembly. The AAGAB psGD:AP1σ3 structure provides one mechanism to achieve such protection. While the SBL loop does not exactly occupy the cargo peptide binding surface, the highly conserved F153, approximating the position of the L(+1) leucine in the cargo peptide, draws AP1σ3 V98 closer to the L0 leucine position, effectively flattening the dileucine binding pocket (Figure 3d, 3e). At the other end, the psGD α5 helix and surrounding regions function as a lid, further preventing cargo binding to this region.

On the psGD side, the most surprising finding from the AAGAB psGD:AP1σ3 crystal structure is the novel interface (Figure 3a-c, 3g, Suppl. Figure 3e). In contrast to the switch I and switch II regions that are regulated by guanine nucleotide binding states, AAGAB psGD employs the opposite surface to engage the σ subunits. Two loops in AAGAB psGD protrude like a pincer to grab the σ subunit (Figure 3a-c). The highly acidic SBL loop docks onto the positively charged surface on AP1σ3 surface (Figure 3a, Suppl. Figure 1a), complementing the σ subunit both in shape and in chemistry. E155, the acidic residue at the center of the SBL loop interface (Figure 3b), is one of the most highly conserved residues in AAGAB proteins across species ^18^. The shorter “jaw” of the pincer, the short turn between β2 and β3, constitutes another set of most highly conserved residues, including K52 and Y53 (Figure 3c). The conservative nature of the interfacial residues, together with our structural, biochemical, and cellular results, argues unambiguously for the novel interface. The discovery is consistent with the pseudoGTPase nature of AAGAB psGD. The psGD does not bind to or hydrolyze GTP, therefore, its interaction is independent of the region equivalent to the nucleotide binding pocket or the switch regions.

Arf1, the small GTPase essential for AP1-mediated membrane trafficking, binds to the β subunit in fully assembled AP1 complex using its switch I/II regions. Crystal structure reveals that Arf1 utilizes a second surface distal to the switch I/II region to weakly engage the γ subunit in another AP1 complex, thus bridging the two AP1 complexes ^44^. Interestingly, the distal region of Arf1 consists of the α4-β6-α5 elements, partially overlapping with the psGD:AP1σ3 interface. It is tempting to speculate that since psGD lacks the canonical switch I/II binding surface, the distal region becomes the major protein-protein interaction surface and evolves to achieve high affinity and specificity in binding.

Two missense mutations have been reported on the pseudoGTPase domain of AAGAB, E144K and V139I ^19,43^. Though the two mutants were comparable in their ability to interact with the σ subunits in in vitro coexpression studies, their thermostability is lower than the wild-type protein (Suppl. Figure 4a-d). In cells, such reduced stability may play a more substantial role in AAGAB dysfunction. Indeed, in zebrafish, the V147I mutant (equivalent to human V139I) fails to ameliorate the larval death phenotype and swimming defects ^17^. In addition, the V139I mutation may impact an AAGAB function independent of σ subunit interaction.

Pseudoenzymes are proteins or protein domains that assume the folds of an active enzyme but are catalytically deficient. Pseudoenzymes exist in multiple enzyme families, including pseudokinases, pseudophosphatases, pseudoproteases, pseudo ubiquitin conjugation and modifying enzymes, and many more ^45,46^. Biologically, pseudoenzymes have been found in all kingdoms of life. It is estimated the 10-15% of the proteome is comprised of pseuduenzymes ^45^. Functions of pseudoenzymes span from allosteric regulation of an active enzyme partner, serving as scaffolds for signaling, to sequestration of substrates. Among pseudoenzymes, pseudoGTPases are less studied. Our study identified AAGAB G domain as a new member of the pseudoGTPase family. Furthermore, we discovered that the pseudoGTPase domain of AAGAB functions as a stabilizer and protector of the AP1 and AP2 σ subunits through direct protein-protein interaction. The interaction is achieved through a surface distal to the conventional switch I and switch II regions regulated by guanine nucleotide binding. The knowledge garnered from investigating AAGAB psGD structure and function may lead to new strategies in protein engineering and targeted therapeutics.

## Methods

### Cloning, coexpression and purification

Full-length human AP1S3 (Uniprot Q96PC3-1) was cloned into pET26b vector (EMD Biosciences) between *Nde*I and *Xho*I sites with a fused His_6_ tag at the C-terminus. The tagless AAGAB psGD domain (residues 2-177) was cloned between *Nde*I and *Xho*I sites at the second multiple cloning site in pETDuet-1 vector (EMD Biosciences). Both constructs were confirmed by DNA sequencing. Equal amounts of these two plasmids were mixed first and used to transform BL21 (DE3) *E. coli* cells selected by Agar plate containing both Kanamycin and Ampicillin. The clones containing both plasmids were overexpressed in LB medium. The cells were induced by 0.4 mM IPTG when OD600 reached a value of ∼ 0.6 and were cultured overnight at 20 °C in an incubator shaker. The cell extract was prepared by sonication in buffer A (50 mM HEPES pH 7.5, 150 mM NaCl, 25 mM imidazole, and 5 mM β-mercaptoethanol). After centrifugation, the supernatant was loaded onto a Ni-affinity column, which was then extensively washed with buffer A and eluted with buffer B (50 mM HEPES pH 7.5, 500 mM NaCl, and 300 mM imidazole pH 8.0). The peak fractions were further purified by size exclusion column chromatography using a Hiload 16/60 Superdex-200 prep grade column (GE healthcare) in buffer C (50 mM HEPES pH 7.5, 150 mM NaCl, and 5 mM β-mercaptoethanol).

Protein purity was assessed by SDS-PAGE. The peak fractions containing pure binary complex protein were pooled and concentrated using 10 kDa cut-off centrifugal concentrator (EMD Millipore).

Wild-type AAGAB psGD (aa 2-177) was cloned into the first multiple cloning site of a SUMO-pRSFDuet-1 vector between the *BamH*I and *Sal*I sites. The SUMO-pRSFDuet-1 vector is modified from pRSFDuet-1 and bears an in-frame N-terminal His_6_-SUMO tag. All AAGAB psGD domain mutants discussed in this study were generated by a two-step PCR-based overlap extension method, cloned in the same way as the wild-type pGD, and confirmed by DNA sequencing. The constructs were overexpressed in *E. coli* strain BL21 (DE3) and the cell extracts were prepared in the same way as the binary complex. Briefly, the proteins were first purified by a Ni-affinity column, followed by SUMO protease Ulp1 treatment overnight at 4 °C to cleave the His_6_-SUMO tag. The His_6_- SUMO tag was removed via a second Ni-affinity column. The tagless proteins, unbound to Ni-affinity column, were concentrated by HiTrap Q column (GE Healthcare). The concentrated proteins were finally purified by gel filtration chromatography using Hiload 16/60 Superdex 200 prep grade column (GE healthcare) in buffer C and concentrated in the same way as the binary complex. All purified proteins were immediately frozen in liquid nitrogen and stored at -80 °C.

### Crystallization and data collection

The purified human AAGAB psGD:AP1σ3 binary complex crystals were grown using the sitting-drop vapor-diffusion method by mixing the protein with an equal volume of reservoir solution containing 0.1 M Bis-Tris pH 6.5, 25% PEG3350 at 16 °C. The standalone AAGAB psGD^E144K^ crystals were grown using the hanging-drop vapor-diffusion method by mixing the protein with an equal volume of reservoir solution containing 0.1 M Bis-Tris pH 5.5 and 29% PEG3350 at 16 °C. The crystals started to show up after 3 days and reached full size within a week. For data collection, all crystals were flash frozen in the oil perfluoropolyether (Hampton Research) as the cryo- protectant. Diffraction datasets were collected on AMX and FMX beamlines at the National Synchrotron Light Source II (NSLS-II) and 24-ID-E beamline at the Advanced Photon Source (APS).

### Structure determination and refinement

The dataset of AAGAB psGD:AP1σ3 was indexed, integrated, and scaled using the XDS program package ^47^. The AAGAB psGD:AP1σ3 crystal belongs to orthorhombic space group P2_1_2_1_2_1_ (a = 50.294 Å, b = 61.608 Å, c = 92.854 Å, α = β = γ =90°) and contains one binary complex per asymmetric unit. The dataset of AAGAB psGD^E144K^ was processed with the HKL2000 suite ^48^. The AAGAB psGD^E144K^ crystal belongs to the orthorhombic space group P2_1_2_1_2_1_ (a = 39.268 Å, b = 94.349 Å, c = 102.796 Å, α = β = γ = 90°) and contains two molecules per asymmetric unit.

The initial phase for AAGAB psGD: AP1σ3 was obtained by molecular replacement with Phenix program Phaser ^49,50^ using AP1σ1 (∼80% identical with AP1σ3) from the AP1 core structure (PDB ID, 1W63) as a search model. The initial resultant map showed great densities for AP1σ3 but not psGD. After first round refinement and adding water molecules, the densities for psGD interfacial residues became visible. The resultant map was then subjected to the AutoBuild program in Phenix for psGD building.

Subsequent iterations of manual building and refinement were conducted in Coot ^51,52^ and Phenix ^50^ to the final R_work_/R_free_ = 18.3/21.1%. The final model of AAGAB psGD:AP1σ3 contains residues 4-30, 40-177 of AAGAB psGD and residues 1-123 of AP1σ3 and 324 water molecules.

The structure of free AAGAB psGD^E144K^ was solved by molecular replacement using the AAGAB psGD model from the complex structure as the search model in Phaser ^49,50^.

Iterations of manual building/adjustment and refinement were conducted in WinCoot ^51,52^ and Phenix ^49,50^ to the final R_work_/R_free_ = 18.8/20.7%. The final model contains residues 2-144 and 156-177 of AAGAB psGD.

### GTPase assay

For measuring the GTPase activity, the malachite green phosphate detection kit (Sigma, Cat No. MAK307) was used. 500 nM AAGAB GD or 500 nM GBP2FL protein was mixed with 200 µM GTP (Sigma, Cat No. G8877) in a reaction buffer containing 20 mM Tris-HCl pH 8.0, 150 mM NaCl, 8 mM MgCl_2_ and 1 mM EDTA at room temperature. Aliquots were taken out at indicated time points up to 60 min and the GTP hydrolysis reaction was stopped by 25 mM EDTA. Reagent A and Reagent B from the kit were mixed in 100:1 ratio. 20 μL of the prepared reagent was mixed with 80 μL of EDTA-stopped reaction mixture, added to 96 well clear plate, and incubated at room temperature for 30 min for malachite green color change to develop. Absorbance was read in the microplate reader (SpectraMax iD5, Molecular Devices) at 620 nm.

### Guanine nucleotide binding assay

The 2’/3’-O-(N-methylanthraniloyl (mant-)) nucleotides mant-GDP and mant-GTPγS (Jena Bioscience) were used as fluorescent probes for determining the binding affinity of AAGAB psGD. GBP2GD and SUMO were included as positive and negative controls, respectively. Different concentrations of proteins were incubated with mant-nucleotides in the binding buffer containing 25 mM Tris-HCl pH 8.0, 150 mM NaCl, 2 mM MgCl_2_, and 5mM β-ME at 25 °C. The mant-nucleotides were excited at 355 nm and the fluorescence was measured at 448 nm. The binding of the mant-nucleotides to the proteins was monitored as increase in fluorescence using a microplate reader (SpectraMax iD5, Molecular Devices). The increase in fluorescence was plotted against the protein concentrations and the dissociation constant was obtained by fitting the data to one-site specific binding model in GraphPad Prism 7.0 (GraphPad Software, San Diego, CA).

### Size-exchange chromatography coupled multiple-angle light scattering (SEC-MALS)

AAGAB psGD:AP1σ3 binary complex was injected into a Superdex 200 Increase 10/300 size-exclusion column equilibrated in a buffer containing 25 mM HEPES pH 7.5 and 150 mM NaCl. The column was coupled with a multi-angle light scattering detector (DAWN HELEOS II, Wyatt Technology) and a refractometer (Optilab T-rEX, Wyatt Technology). Data were collected every 0.5 sec at a flow rate of 0.5 mL/min at room temperature. Data processing was carried out using the program ASTRA 7.3 (Wyatt Technology). Molar mass and mass distribution of the AAGAB psGD:AP1σ3 binary complex were calculated and reported by ASTRA 7.3.

### Pull-down assay using coexpressed recombinant proteins

GST-AP1σ3 and GST-AP2σ1 were expressed individually or in combination with His_6_- SUMO-psGD WT or Y2R, A168R, or DFE3R plasmids in *E. coli* BL21(DE3) cells. Cells were cultured in 500 mL LB medium with corresponding antibiotics. After harvesting, cells were lysed and centrifuged following the same protocol described above. The cleared cell lysates were equally split for either GST or nickel affinity pull-down. For GST affinity pull-down, the cleared cell lysate was loaded onto a gravity column with 1 mL bed volume of Glutathione agarose resin (Gold Biotechnology, #G-250-5). The resin was extensively washed, and bound proteins were eluted with a GST elution buffer (50 mM Tris-HCl, pH 8.0, 150 mM NaCl, and 30 mM L-Glutathione reduced) in 1 mL fractions. For nickel affinity pull-down, the cleared cell lysate was loaded onto a gravity column with 0.5 mL bed volume of Ni-NTA resin (Qiagen, #30210). The resin was extensively washed, and bound proteins were eluted with a Ni-NTA elution buffer (50 mM Tris-HCl, pH 8.0, 150 mM NaCl, and 300 mM imidazole) in 0.5 mL fractions. All elution fractions were analyzed by SDS-PAGE.

### Immunoblotting and immunoprecipitation

To detect proteins in whole cell lysates, cells grown in 24-well plates were lysed in the SDS protein buffer. Protein samples were resolved on 8% Bis-Tris SDS-PAGE and proteins were detected using primary antibodies and horseradish peroxidase (HRP)- conjugated secondary antibodies. Primary antibody used in this work included monoclonal anti-FLAG antibodies (Millipore-Sigma, #F1804) at a final concentration of 1 μg/mL, polyclonal anti-AP1γ antibodies (Bethyl, #A304-771A) at a final concentration of 1 μg/mL, polyclonal anti-AP2α antibodies (BD Biosciences, #610502) at a final concentration of 1 μg/mL, polyclonal anti-AP1σ antibodies (Bethyl, #A305-396A) at a final concentration of 1 μg/mL, monoclonal anti-AP2σ antibodies (Abcam, #ab92380) at a final concentration of 1 µg/mL, and monoclonal anti-α-tubulin antibodies (DSHB, #12G10) at a final concentration of 43 ng/mL.

In immunoprecipitation (IP) experiments, cells were lysed in IP buffer [25 mM HEPES (pH 7.4), 138 mM NaCl, 10 mM Na_3_PO_4_, 2.7 mM KCl, 0.5% CHAPS, 1 mM DTT, and a protease inhibitor cocktail]. Transiently expressed 3xFLAG-AAGAB was precipitated using Protein A/G agarose beads and anti-FLAG M2 antibodies (Millipore-Sigma, #F1804) at a final concentration of 0.5 μg/mL. Proteins in the immunoprecipitates were detected using immunoblotting.

### Flow cytometry

HeLa cells were maintained in Dulbecco’s Modified Eagle Medium (DMEM) supplemented with 10% FB essence (FBE; Seradigm, #3100-500) and penicillin-streptomycin (Millipore-Sigma, #P4333). To stain surface TfR, HeLa cells were washed three times with the KRH buffer (121 mM NaCl, 4.9 mM KCl, 1.2 mM MgSO_4_, 0.33 mM CaCl_2_, and 12 mM HEPES, pH 7.0). Cells were then chilled on ice and stained with monoclonal anti-TfR antibodies (DSHB, #G1/221/12) at a final concentration of 0.1 μg/mL and APC-conjugated secondary antibodies (Thermo Fisher Scientific, #17-4015- 82) at a final concentration of 0.8 μg/mL ^53^. After dissociation from plates using Accutase (Innovative Cell Technologies, #AT104), APC fluorescence of the cells was measured on a CyAn ADP analyzer (Beckman Coulter). Mean APC fluorescence of cells expressing WT and mutant AAGAB proteins was normalized to the control sample in which an empty vector was transfected into *AAGAB* KO cells. Data from populations of ∼5,000 cells were analyzed using the FlowJo software (FlowJo, LLC, v10) based on experiments run in biological triplicates.

### Immunostaining and Imaging

Cells grown on microscope cover glasses (VWR, #89015-725) were washed three times with KRH buffer and fixed using 4% paraformaldehyde. Surface TfR was stained using monoclonal anti-TfR antibodies (DSHB, #G1/221/12) at a final concentration of 0.1 μg/mL and Alexa Fluor 568-conjugated secondary antibodies (Thermo Fisher Scientific, #A11004) at a final concentration of 1 μg/mL. The nuclei were stained with Hoechst 33342 (Thermo Fisher Scientific, #H3570) at a final concentration of 10 μg/mL ^53^.

Images were captured using a 100× oil immersion objective on a Nikon A1 Laser Scanning confocal microscope and processed using FIJI software.

## Data availability

The atomic coordinates and structural factors of AAGAB psGD^E144K^ and psGD:AP1σ3 complex have been deposited in the Protein Data Bank with the accession codes 9DDS (AAGAB psGD^E144K^) and 9DDT (AAGAB psGD:AP1σ3 complex).

## Acknowledgments

We thank Drs. James Hurley and Juan Bonifacino for reagents or advice. We thank Dr. T. “Soma” Somasundaram and beamline scientists at APS 22-ID and 22-BM and NSLSII AMX and FMX (17-ID-1 and 17-ID-2) for assistance in X-ray diffraction data collection. We thank Dr. Gwimoon Seo for competent cells and 3C and Ulp1 proteases. We thank Dr. Peter Randolph for his assistance in SEC-MALS data analysis. This work was supported by the National Institutes of Health grants GM138685 (Q.Y.), AI146330 (Q.Y.), GM126960 (J.S.).

## Author Contributions

Y.T., S.L., J.S., and Q.Y. conceived the initial experimental plan. Y.T., R.Y., and B.W. expressed and purified the proteins. B.W., Y.T. and Q.Y. carried out crystallization and structural determination and refinement. R.Y. performed pull down assay, nucleotide binding assay, and SEC-MALS analysis. S.R. carried out the GTPase activity assay. C.W. and J.W. performed all cell-based assays. B.W., Y.T., S.L., J.S., and Q.Y. drafted the manuscript. All authors edited the manuscript.

