## Supplementary Method for "Structural basis of pseudoGTPase-mediated protein-protein interactions"

*Thermal shift assay*

AAGAB psGD WT, E144K, and V139I mutants were mixed with the Protein Thermal Shift Dye™ from the Protein Thermal Shift™ Dye Kit (Applied Biosystems, Waltham, MA) to a final reaction volume of 20 µl. Three final protein concentrations were tested: 5 µM, 10 µM, and 15 µM. Protein-dye mixes were prepared following the manufacturer's instructions in PCR strips. They were heated in a temperature gradient of 25 - 95 °C with 0.5 °C increment every 15 seconds in a CFX96 real-time PCR system (Bio-Rad Laboratories, Hercules, CA). Fluorescence signal was recorded in triplets with ROX as the fluorophore. The data were fitted to the Boltzmann equation to obtain melting temperature using GraphPad Prism 7.0 (GraphPad Software, San Diego, CA).
