## Supplementary Figures 1-5 for "Structural basis of pseudoGTPase-mediated protein-protein interactions"

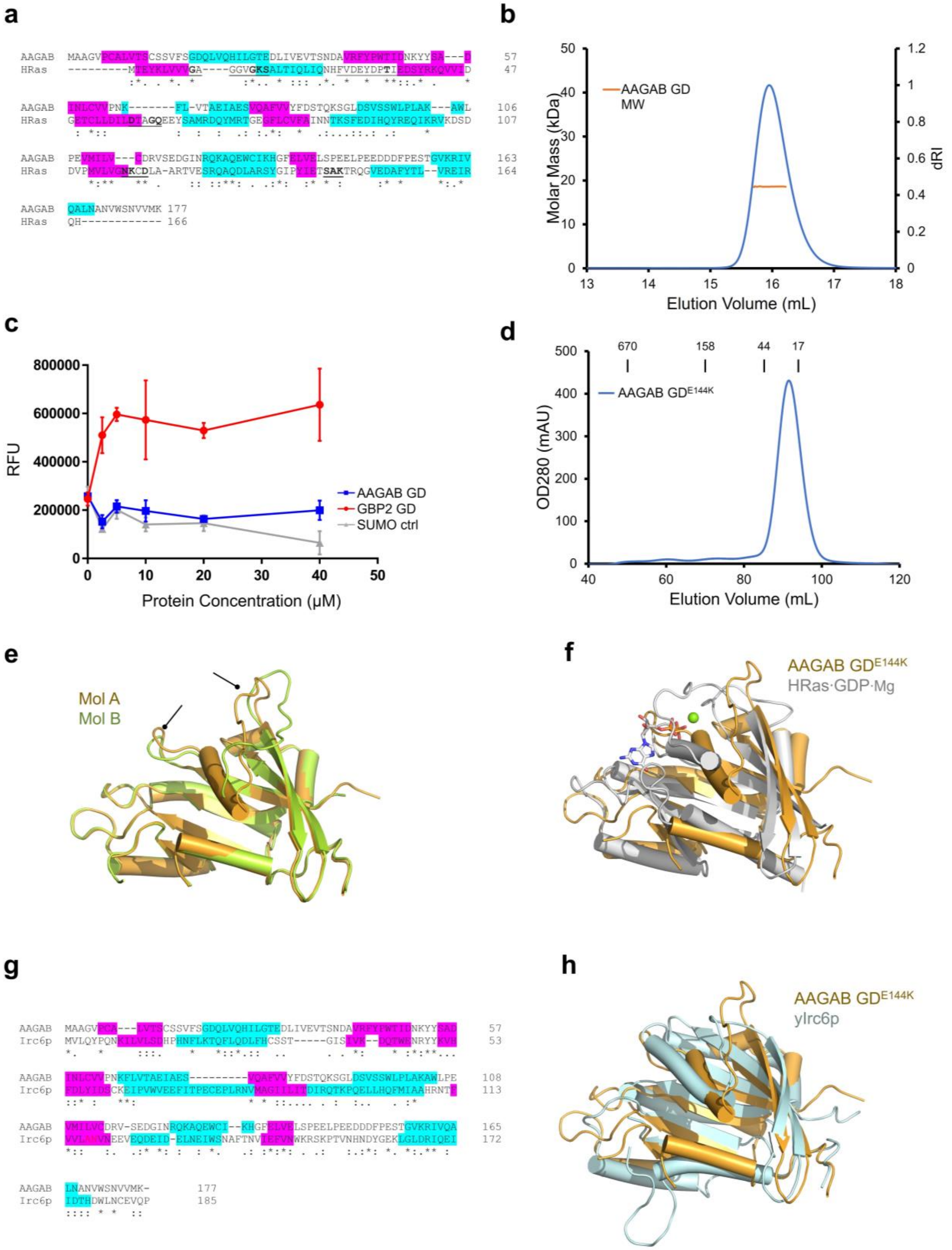
a
AAGAB MAAGVPCALVTSCSSVFSGDQLVQHILGTEDLIVEVTSNDAVRFYPWTIDNKYYSA---D 57
HRas ---------MTEYKLVVVGA----GGVGKSALTIQLIQNHFVDEYDPTIEDSYRKQVVID 47
AAGAB INLCVVPNK-------FL-VTAEIAESVQAFVVYFDSTQKSGLDSVSSWLPLAK---AWL 106
HRas GETCLLDILDTAGQEEYSAMRDQYMRTGEGFLCVFAINNTKSFEDIHQYREQIKRVKDSD 107
AAGAB PEVMILV---CDRVSEDGINRQKAQEWCIKHGFELVELSPEELPEEDDDFPESTGVKRIV 163
HRas DVPMVLVGNKCDLA-ARTVESRQAQDLARSYGIPYIETSAKTRQGVEDAFYTL--VREIR 164
AAGAB QALNANVWSNVVMK 177
HRas QH------------ 166
b
AAGAB GD
MW
Molar Mass (kDa)
dRI
Elution Volume (mL)
c
RFU
AAGAB GD
GBP2 GD
SUMO ctrl
Protein Concentration (μM)
d
670
158
44
17
AAGAB GDE144K
OD280 (mAU)
Elution Volume (mL)
e
Mol A
Mol B
f
AAGAB GDE144K
HRas·GDP·Mg
g
AAGAB MAAGVPCA---LVTSCSSVFSGDQLVQHILGTEDLIVEVTSNDAVRFYPWTIDNKYYSAD 57
Irc6p MVLQYPQNKILVLSDHPHNFLKTQFLQDLFHCSST-----GISIVK--DQTWENRYYKVH 53
AAGAB INLCVVPNKFLVTAEIAES---------VQAFVVYFDSTQKSGLDSVSSWLPLAKAWLPE 108
Irc6p FDLYIDSCKEIPVWVEEFITPECEPLRNVMAGIILITDIRQTKPQELLHQFMIAAHRNTF 113
AAGAB VMILVCDRV-SEDGINRQKAQEWCI--KHGFELVELSPEELPEEDDDFPESTGVKRIVQA 165
Irc6p VVLANVNEEVEQDEID-ELNEIWSNAFTNVIEFVNWKRSKPTVNHNDYGEKLGLDRIQEI 172
AAGAB LNANVWSNVVMK- 177
Irc6p IDTHDWLNCEVQP 185
h
AAGAB GDE144K
yIrc6p

**Supplementary Figure 1 AAGAB N-terminal region is a pseudoGTPase domain**

(a) Sequence alignment of human AAGAB N-terminal region with human HRas. Identical and similar residues are marked by *, : and .. The five G motifs in HRas are underlined, and consensus residues are in bold. α helices and β strands are highlighted in cyan and pink, respectively. (b) Size-exclusion chromatography (SEC) coupled multi-angle light scattering (SEC-MALS) profile of AAGAB G domain (GD). Blue trace: refractive index (RI) of AAGAB normalized to 1; orange trace: calculated MW across the AAGAB GD peak. (c) AAGAB GD (blue), GBP2 GD (red), and SUMO (gray) are mixed with mant-GDP and changes in mant-GDP fluorescence are monitored. (d) SEC profile of AAGAB GD E144K mutant from a HiLoad 16/600 column. Protein standards with known molecular weight are marked at the top. (e) Superposition of the two $GD^{E144K}$ molecules mol A (gold) and mol B (yellow-green) in the asymmetric subunit. Arrows point to the two loops with different conformations. (f) Structure superposition of AAGAB psGD (gold) with HRas·GDP·Mg (gray, PDB ID 4Q21). (g) Sequence alignment of human AAGAB N-terminal region with yeast Irc6p. Identical and similar residues are marked by *, : and .. α helices and β strands are highlighted in cyan and pink, respectively. (h) Structure superposition of AAGAB psGD (gold) with yeast Irc6p (pale cyan, PDB ID 3UC9).

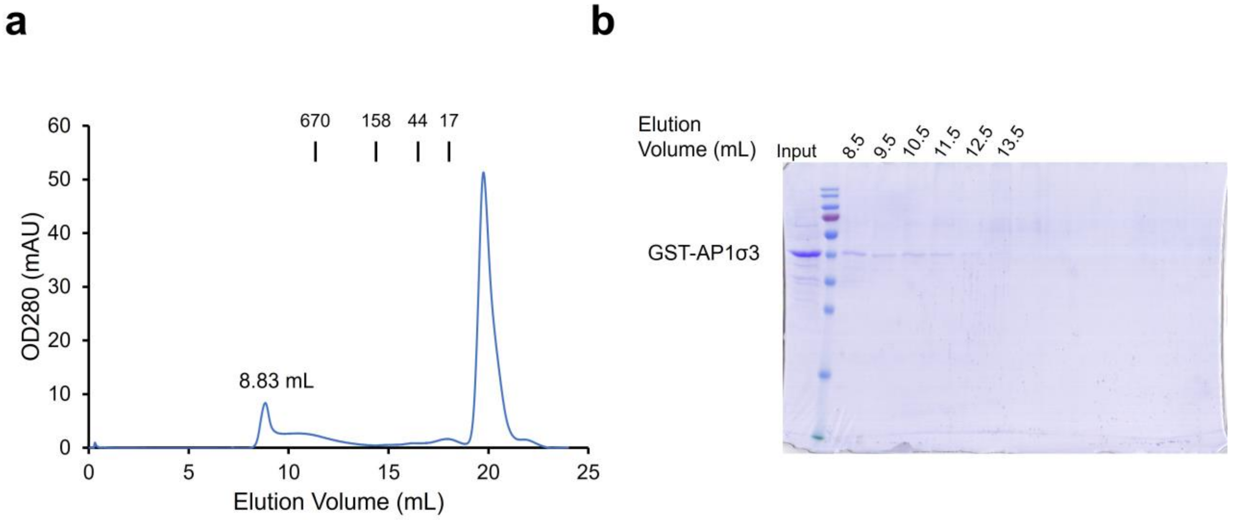


**Supplementary Figure 2 AAGAB psGD assists in σ subunit folding**

(a) Size exclusion chromatography profile of GST-AP1σ3 on a Superdex 200 10/300 column. Protein standards with known molecular weight are marked at the top. (b) SDS-PAGE of elution fractions in (a).

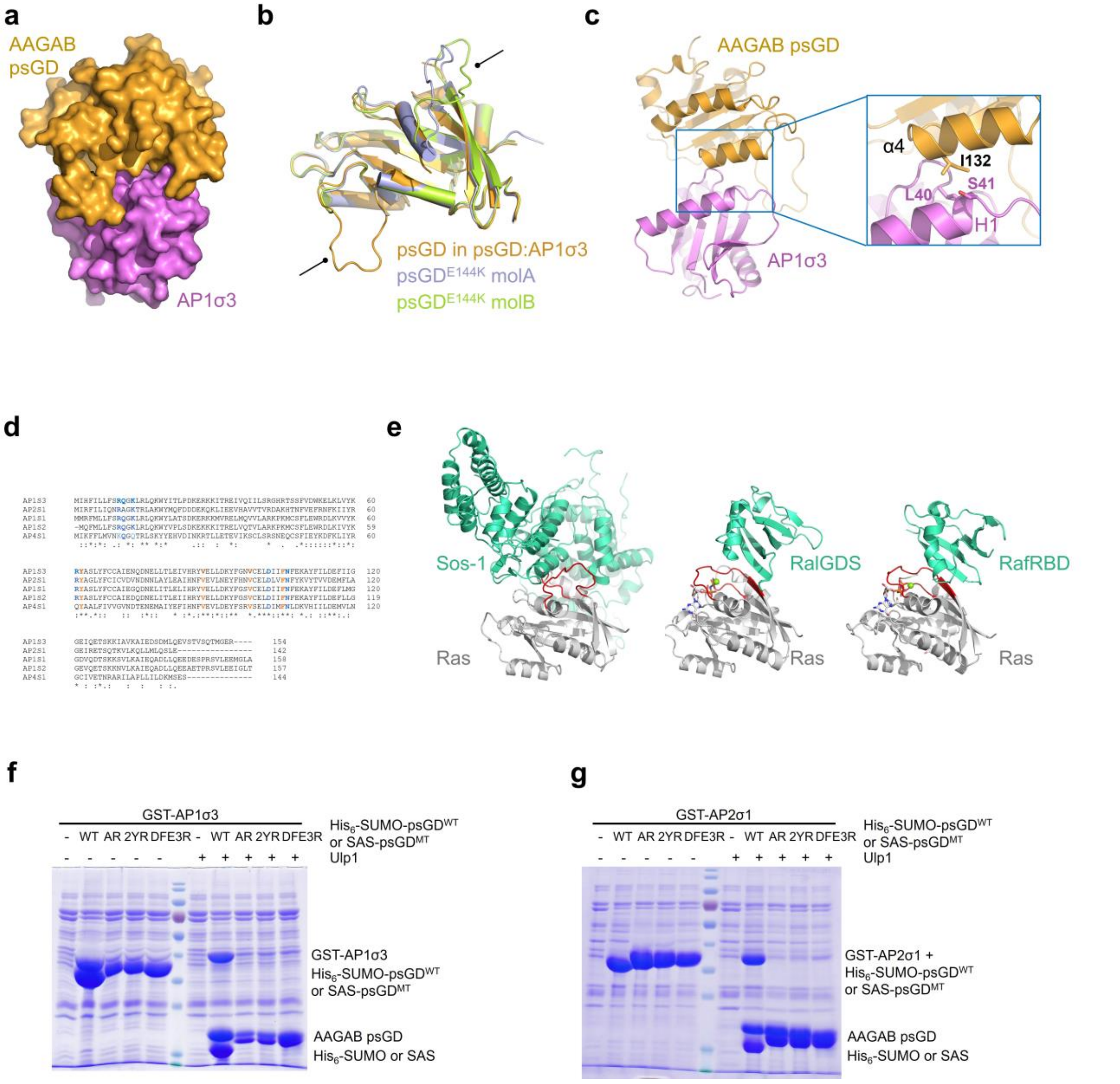


**Supplementary Figure 3 AAGAB psGD utilizes a unique interface to interact with sigma adaptins**

(a) Surface presentation of AAGAB psGD:AP1σ3 complex oriented the same as in Figure 3a left panel. psGD and AP1σ3 are colored gold and violet, respectively. (b) Superposition of AAGAB psGD in psGD:AP1σ3 complex (gold), AAGAB $GD^{E144K}$ molecule A (light blue), and AAGAB $GD^{E144K}$ molecule B (lemon). (c) AAGAB psGD:AP1σ3 interaction between psGD α4 and AP1σ3 H1 (left panel). The presentation in (c) is rotated ~ 160 ° along y axis from in (a). The key residues psGD I132 and AP1σ3

L40 and S41 are shown as sticks in the zoom-in view (right panel). (d) Sequence alignment of human AP1σ3, AP2σ1, AP1σ1, AP1σ2, and AP4σ1. UniProt ID: AP1σ3: Q96PC3; AP2σ1: P53680; AP1σ1: P61966; AP1σ2: P56377; AP4σ1: Q9Y587. Residues interacting with psGD are in bold and color blue (hydrophilic residues) or orange (hydrophobic residues). * identical residues; : similar residues. (e) Structures of HRas:Sos-1 (PDB ID: 1BKD), HRas: RalGDS (PDB ID: 1LFD), and HRas:RafRBD (PDB ID: 4G0N). The orientations of HRas molecules are very similar to that in Figure 3g. HRas molecules are colored gray, while Sos-1, RalGDS, and RafRBD are colored in greencyan. Switch I and II regions of HRas are highlighted in red. (f) Nickel pull-down gel of GST-AP1σ3 co-expressed with $His_6$-SUMO tagged WT or SAS tagged mutant AAGAB psGD. AR: A168R; 2YR: Y53R/Y54R; DEF3R: D151R/F153R/E155R. Ulp1 protease treatment is used to confirm the identity of $His_6$-SUMO-AAGAB $psGD^{WT}$ and SAS-AAGAB psGD mutants. (g) Nickel pull-down gel of GST-AP2σ1 co-expressed with $His_6$-SUMO tagged WT or SAS tagged mutant AAGAB psGD. AR: A168R; 2YR: Y53R/Y54R; DEF3R: D151R/F153R/E155R. Ulp1 protease treatment is used to confirm the identity of $His_6$-SUMO-AAGAB $psGD^{WT}$ and SAS-AAGAB psGD mutants. Note that the molecular weights of GST-AP2σ1 and $His_6$-SUMO-AAGAB psGD are too close to separate on SDS-PAGE.

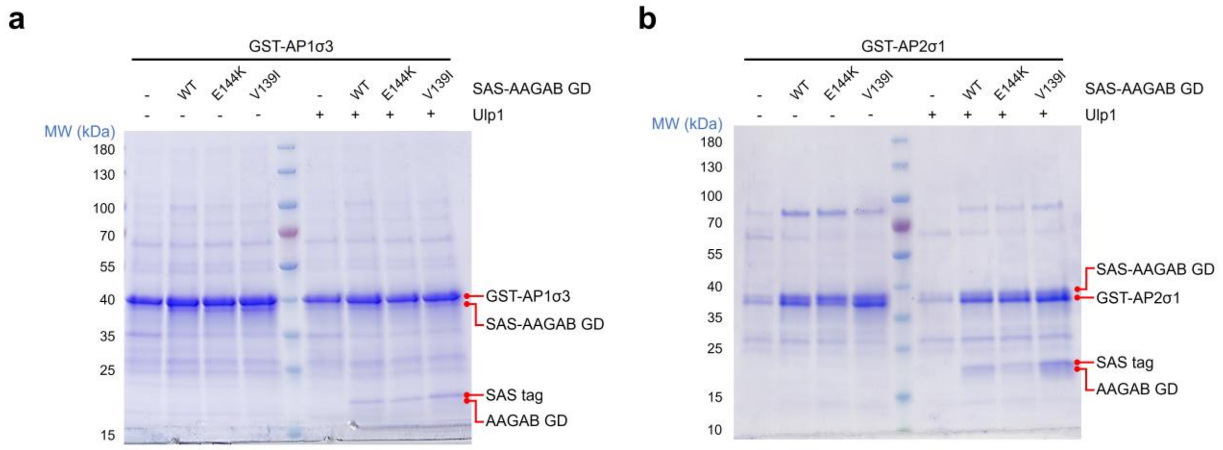


c

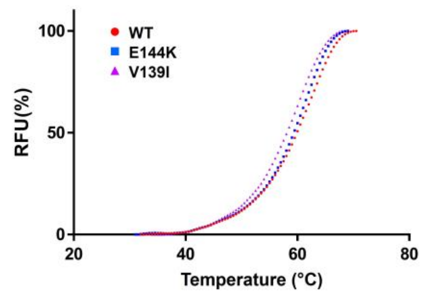


d

| Protein Concentration | WT | E144K | V139I |
|---|---|---|---|
| 5 μM | 61.2 ± 0.3 | 60.7 ± 0.3 | 59.1 ± 0.2 |
| 10 μM | 61.3 ± 0.3 | 60.5 ± 0.3 | 59.8 ± 0.3 |
| 15 μM | 60.9 ± 0.3 | 60.0 ± 0.3 | 59.3 ± 0.3 |

**Supplementary Figure 4 Characterization of PPKP1-related psGD mutants E144K and V139I**

(a) GST pull-down gel of GST-AP1σ3 co-expressed with SAS tagged AAGAB psGD WT, E144K, or V139I mutant. Ulp1 protease treatment is used to confirm the identity of SAS-AAGAB psGD. (b) GST pull-down gel of GST-AP2σ1 co-expressed with SAS tagged AAGAB psGD WT, E144K, or V139I mutant. Ulp1 protease treatment is used to confirm the identity of SAS-AAGAB GD. (c) A representative thermostability assay of AAGAB psGD WT, E144K, and V139I at the protein concentration of 5 μM. The fluorescence values are normalized to the maximum value at the highest temperature point for each psGD protein. (d) Summary of the melting temperatures (*T*m) for AAGAB psGD WT, E144K, and V139I at three different protein concentrations.

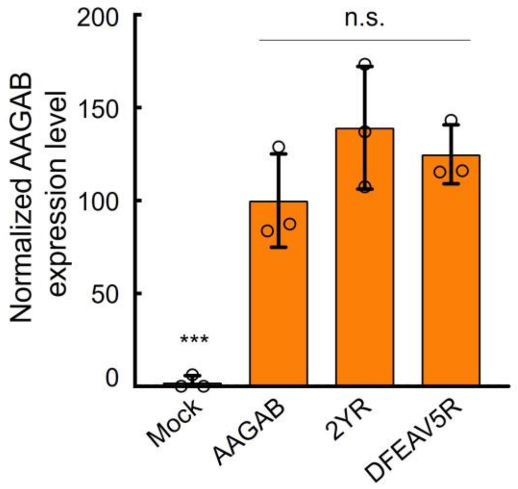


**Supplementary Figure 5 Expression levels of AAGAB WT and psGD mutants in cells**

The expression levels of indicated protein were normalized to WT AAGAB samples. Data are presented as mean±s.d., n=3. n.s., P>0.05. P values were calculated using one-way ANOVA and Dunnett’s multiple comparison test. The representative immunoblots are shown in Figure 3e.
