## Supplementary Table 1 for "Structural basis of pseudoGTPase-mediated protein-protein interactions"

| | AAGAB psGD E144K | psGD:AP1σ3 |
|---|---|---|
| **Data Collection** | | |
| Wavelength (Å) | 0.99228 | 0.97918 |
| Resolution range (Å) | 45.14 – 1.78 (1.844 – 1.78) | 46.43 – 1.68 (1.74 – 1.68) |
| Space group | P 21 21 21 | P 21 21 21 |
| Unit cell (a, b, c, Å)<br>(α, β, γ, °) | 39.268 94.349 102.796<br>90 90 90 | 50.294 61.608 92.854<br>90 90 90 |
| Unique reflections | 34572 (2096) | 33173 (3112) |
| Multiplicity | 9.6 (2.9) | 12.9 (11.9) |
| Completeness (%) | 93 (63) | 98.68 (94.16) |
| Mean I/sigma(I) | 14.4 (0.7) | 18.51 (2.39) |
| Wilson B-factor | 23.70 | 26.20 |
| R-merge | 0.119 (0.841) | 0.07416 (0.8291) |
| R-meas | 0.125 (0.985) | 0.07728 (0.8663) |
| R-pim | 0.037 (0.494) | 0.02136 (0.2458) |
| CC1/2 | 1.004 (0.508) | 0.999 (0.872) |
| CC* | | 1 (0.965) |
| **Refinement** | | |
| Reflections used in refinement | 34542 (2082) | 33173 (3111) |
| Reflections used for R-free | 1980 (119) | 1995 (187) |
| R-work | 0.1883 (0.3501) | 0.1825 (0.2267) |
| R-free | 0.2071 (0.3737) | 0.2113 (0.2546) |
| Number of non-hydrogen atoms | 2942 | 2669 |
| macromolecules | 2569 | 2346 |
| solvent | 373 | 323 |
| Protein residues | 330 | 288 |
| RMS (bonds, Å) | 0.009 | 0.008 |
| RMS (angles, °) | 1.20 | 1.07 |
| Ramachandran favored (%) | 99.07 | 99.29 |
| Ramachandran allowed (%) | 0.93 | 0.71 |
| Ramachandran outliers (%) | 0.00 | 0.00 |
| Rotamer outliers (%) | 1.03 | 0.38 |
| Clashscore | 0.98 | 4.06 |
| Average B-factor | 28.49 | 30.40 |
| macromolecules | 27.49 | 29.44 |
| solvent | 35.34 | 37.44 |
| **Number of TLS groups** | 1 | 1 |

**Table S1.** Crystallography data collection and refinement statistics. Values in parentheses are for the highest resolution shell.
